# Functional consequences of centromeric satellite array asymmetry in female meiosis

**DOI:** 10.64898/2026.05.15.725525

**Authors:** Jun Ma, Deepanshu Kumar, Advait Thaploo, Liya Ma, Michael A. Lampson

## Abstract

Centromeres are epigenetically specified chromosomal sites that support kinetochore assembly and often embedded within large satellite DNA arrays. Recent telomere to telomere genome assemblies have revealed extensive variation in centromeric satellite arrays between chromosomes and between individuals, but the functional significance of this variation remains unclear. To determine how satellite array size influences centromere function, we generated hybrid mouse models in which homologous chromosomes with different array sizes are paired in meiosis I, creating array size asymmetry across each meiotic bivalent. When an extremely small array is paired with a moderate size array, we find that array size asymmetry leads to functional asymmetry in both centromere chromatin and interactions with spindle microtubules, lagging chromosomes in anaphase I, and increased aneuploidy in MII eggs. In contrast, pairing an extremely large array with a moderate array does not lead to functional centromere asymmetry. Together, these results suggest a threshold model in which centromere array size is tolerated across a broad range, but minimal arrays become functionally limiting when paired with larger arrays in meiosis.

## Introduction

Centromeres play a central role in cell division by serving as the chromosomal sites for assembling kinetochores, which mediate interactions between chromosomes and spindle microtubules. In most mammals and many other eukaryotes, centromeres are embedded within long arrays of highly repetitive satellite DNA. Advances in long-read genome assemblies have uncovered extensive variation in the composition, size, and organization of these arrays between chromosomes and between individuals (Miga et al., 2020; Altemose et al., 2022; Logsdon et al., 2024). The functional consequences of this variation remain unclear, however, because centromeres are epigenetically specified by nucleosomes containing a histone H3 variant, CENP-A, suggesting that centromere function may not depend on the underlying DNA sequence (Sundararajan and Straight, 2022; Enriquez and Nechemia, 2025). Moreover, only a small fraction of each satellite array is typically occupied by CENP-A nucleosomes, with the majority of the array lying outside the centromere chromatin (Iwata-Otsubo et al., 2017; Altemose at el., 2022; Carty et al., 2025).

Subtle differences in centromere function may be difficult to detect in mitosis, as sister chromatids should have identical centromere satellite arrays, so interactions with spindle microtubules should be symmetrical for each pair of sisters. In contrast, homologous chromosomes are paired in meiosis I, offering the possibility of different centromere satellite arrays on either side of a meiotic bivalent. In this case, functional differences between arrays on the homologous chromosomes may lead to asymmetric interactions with spindle microtubules, force imbalances, and potentially segregation errors. A study of human trisomy 21 provides initial support for the sensitivity of meiosis I to satellite DNA variation. Analyses of parent-child trios show an association between trisomy in the child and maternal asymmetry in chromosome 21, with one homolog carrying a centromere with minimal satellite array size (Mastrorosa et al., 2025). However, these analyses are limited by small sample sizes and the inability to conduct controlled crosses between parents with different array sizes. Hybrid mouse models provide a tractable framework for such crosses, as inbred strains exhibit substantial variation in centromere satellite array size (Arora et al., 2021; Chaudhry et al., 2025), enabling the generation of bivalents with defined centromere satellite array size asymmetry between homologous chromosomes.

Previous studies using hybrids with the CHPO (ZALENDE) mouse strain, which has centromeric satellite arrays at the extreme lower end of the observed size distribution (Arora et al., 2021), demonstrated that array size asymmetry can produce functional asymmetries in recruitment of centromere proteins and interactions with spindle microtubules (Chmátal et al., 2014; Iwata-Otsubo et al., 2017; Akera et al., 2017, 2019). These studies have been limited, however, by the presence of meiotic trivalents, as CHPO contains multiple Rb fusions, common chromosomal rearrangements that join two telocentric chromosomes at their centromeres to create one metacentric chromosome. Each metacentric chromosome pairs with two homologous telocentric chromosomes in the hybrid, generating trivalents that delay progression through metaphase I and are intrinsically prone to segregation errors. Effects of satellite array size asymmetry on chromosome segregation fidelity are difficult to uncouple from effects of trivalents in CHPO hybrids.

Here, we established hybrid mouse systems spanning the observed centromere satellite array size distribution over two orders of magnitude, using inbred strains with small (CHPO), moderate (C57BL/6 and FVB), and large (PWK) arrays (Figure 1A) (Arora et al., 2021). To avoid complications of meiotic trivalents, we backcrossed CHPO to C57BL/6 to generate strains with telocentric chromosomes with CHPO centromeres but without metacentric chromosomes. Crossing two of these strains produces a hybrid with multiple asymmetric bivalents, in which a small CHPO centromere is paired with a larger C57BL/6 centromere, alongside symmetric bivalents that serve as internal controls. We also generated PWK × FVB hybrids (large × moderate), in which the magnitude of the array size asymmetry is similar to CHPO × C57BL/6 (small × moderate). Using these systems, we show that when satellite arrays are reduced to a minimal size (i.e., CHPO), pairing with a larger array creates functional asymmetry within the bivalent, leading to increased chromosome segregation errors. In contrast, pairing a moderate-sized array with a larger array does not lead to such defects. Together, these findings indicate that centromere satellite array asymmetry promotes meiotic aneuploidy when one homolog falls below a functional array size threshold.

**Figure 1:**
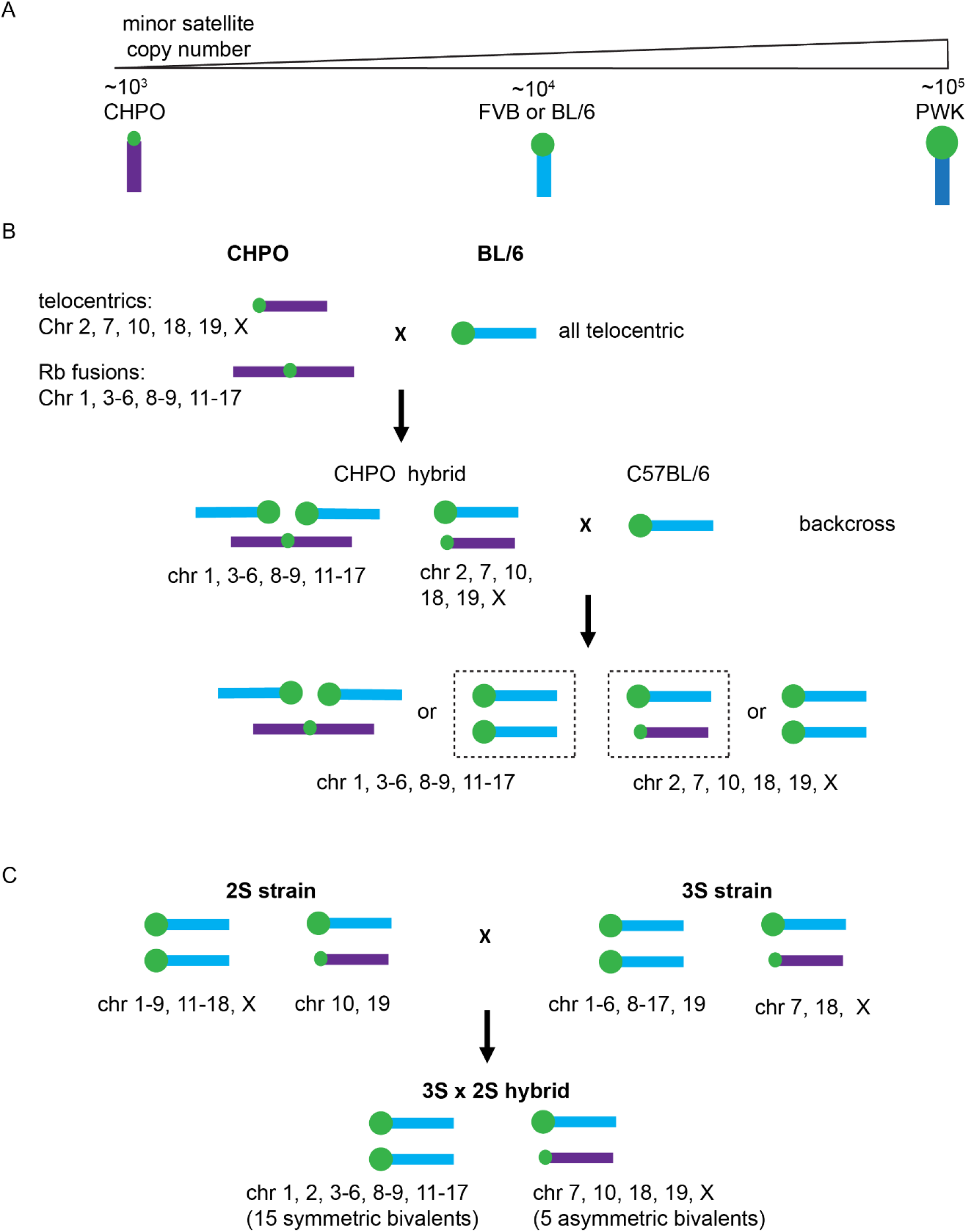
Generation of mouse strains with small centromere satellite arrays without Rb fusion chromosomes. A. Range of centromeric satellite array sizes observed across inbred mouse strains. Sizes of the green circles indicate satellite array sizes, not drawn to scale relative to actual satellite copy number differences. B) Breeding strategy. CHPO mice with smaller centromeric satellite arrays were crossed with C57BL/6 (BL6). The resulting CHPO hybrids were backcrossed to BL6, and progeny were genotyped based on strain-specific SNPs proximal to centromeres to determine the parental origin of each centromere. Progeny in each generation were selected to enrich for telocentric chromosomes with CHPO-derived centromeres and eliminate metacentric Rb fusions. Dashed boxes indicate chromosomes that were enriched by selection. Crossing between progeny generated the 3S strain homozygous for three small centromere arrays (Chr7, 18, and X) and the 2S strain homozygous for two small centromere arrays (Chr10 and 19). Rb fusions are absent in both strains. C) Crossing the 3S and 2S strains generates 3S × 2S hybrid oocytes containing five asymmetric bivalents in meiosis, in which a small centromere satellite array from CHPO is paired with a large array from BL6. The remaining bivalents are symmetric with large arrays.

## Results

### A hybrid system to analyze asymmetries involving small centromere arrays in meiosis

To eliminate metacentric chromosomes from CHPO while maintaining telocentric chromosomes with small centromere satellite arrays, we crossed CHPO with C57BL/6 and subsequently backcrossed to C57BL/6 over multiple generations while selecting for telocentric chromosomes using centromere-linked SNP markers. This strategy generated one strain (3S) homozygous for three CHPO centromeres (Chr 7, 18, and X) and another strain (2S) homozygous for two CHPO centromeres (Chr 10 and 19) (Figure 1B). Crossing these strains produces a hybrid (3S x 2S) containing 5 meiotic bivalents with asymmetric satellite arrays, while the remaining 15 bivalents are symmetric with C57BL/6 centromeres (Figure 1C). To visualize array size asymmetry, we expressed GFP-tagged CENP-B in oocytes from 3S × 2S hybrid females. CENP-B binds a motif (the CENP-B box) in minor satellite DNA, making GFP signal intensity proportional to satellite copy number (Iwata-Otsubo et al., 2017). We observed large differences in GFP intensity across asymmetric bivalents, consistent with the unequal satellite array sizes (Figure 2A).

**Figure 2:**
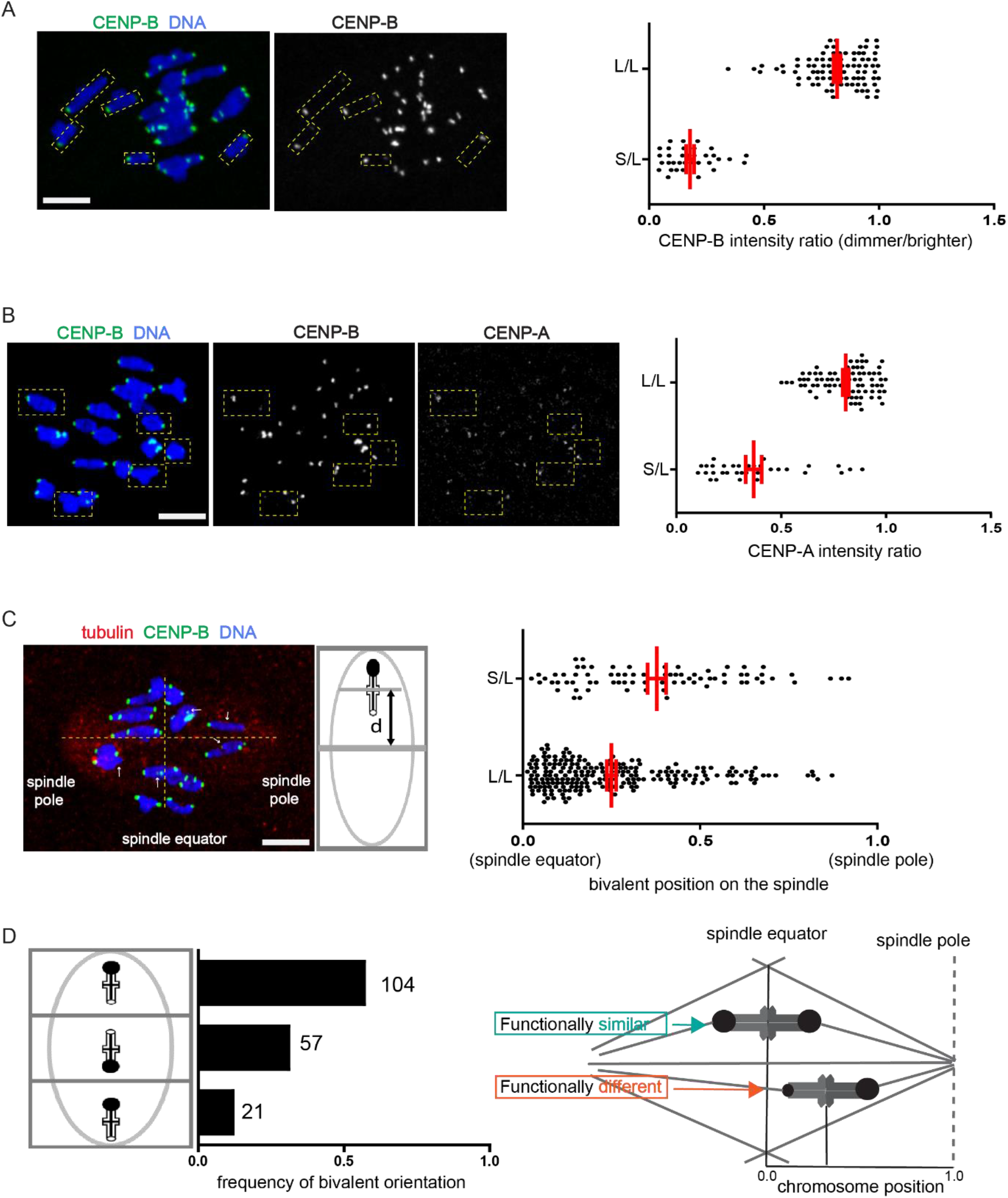
Asymmetric bivalents in 3S × 2S hybrids exhibit CENP-A asymmetry and off-center position on the MI spindle. Oocytes from 3S × 2S hybrids expressing CENP-B–EGFP were fixed at metaphase I and stained for CENP-A or tubulin as indicated. Based on CENP-B intensities, individual bivalents were categorized as asymmetric with one small array and one large array (S/L, yellow boxes in images) or symmetric with two large arrays (L/L). Images are maximum-intensity z-projections. Scale bars 10μm. (A) The ratio of CENP-B intensities between centromeres of paired homologous chromosomes were calculated as dimmer/brighter CENP-B signals. Each dot represents a single bivalent; n = 34 S/L or 86 L/L, pooled from 10 oocytes from 2 independent experiments. (B) The ratio of CENP-A intensities between centromeres of paired homologous chromosomes was calculated as small/large for asymmetric bivalents (S/L, n=33), or as dimmer/brighter CENP-A signals for symmetric bivalents (L/L, n=77). Each dot represents a single bivalent; data are pooled from 10 oocytes from 2 independent experiments. (C) Bivalent positions along the spindle axis (horizonal yellow line) were measured relative to the spindle equator (vertical yellow line). Distance from the spindle equator to the pole is defined as 1 for each cell to normalize for variation in spindle size. Each dot represents a single bivalent; n = 75 S/L or 224 L/L, pooled from 25 oocytes from 2 independent experiments. (D) The orientation of each asymmetric bivalent was categorized as off-center with the larger array closer to the spindle pole (top bar), centered near the spindle equator (middle bar), or off-center with the smaller array closer the spindle pole (bottom bar). Off-center position indicates functional asymmetry between the centromeres of the paired homologous chromosomes.

### Functional consequences of asymmetry with small centromere arrays

To test whether the functional asymmetries previously observed in CHPO hybrid oocytes are maintained in the 3S × 2S hybrid, we measured CENP-A levels and chromosome position on the MI spindle as readouts for centromere chromatin and interactions with spindle microtubules, respectively (Chmatal et al., 2014, Iwata-Otsubo et al., 2017, Akera 2019, Kumon 2021). Within each of the 5 asymmetric bivalents identified based on CENP-B-GFP, the centromere with the smaller satellite array showed reduced levels of CENP-A, whereas CENP-A was similar across each of the 15 symmetric bivalents (Figure 2B). Asymmetric bivalents were also frequently positioned away from the spindle equator, closer to the pole facing the centromere with the larger array, indicating differential interactions with spindle microtubules between centromeres with different satellite array sizes (Figure 2C, D). In contrast, symmetric bivalents were positioned near the equator. These functional asymmetries in centromere chromatin and microtubule interactions are consistent with our previous findings in CHPO hybrid oocytes (Chmátal et al., 2014; Iwata-Otsubo et al., 2017).

To assess chromosome segregation fidelity in 3S × 2S hybrid oocytes compared to 2S control oocytes, we performed live imaging of anaphase I with a DNA-binding dye (Figure 3A). We find frequent segregation errors in hybrids during anaphase I, most commonly in the form of lagging chromosomes. We also find more severe defects in which multiple chromosomes remained clustered or stretched within the spindle and did not resolve into two distinct groups, leading to gross segregation failure. Both lagging chromosomes and gross segregation failures are more frequent in hybrid oocytes compared to controls.

**Figure 3:**
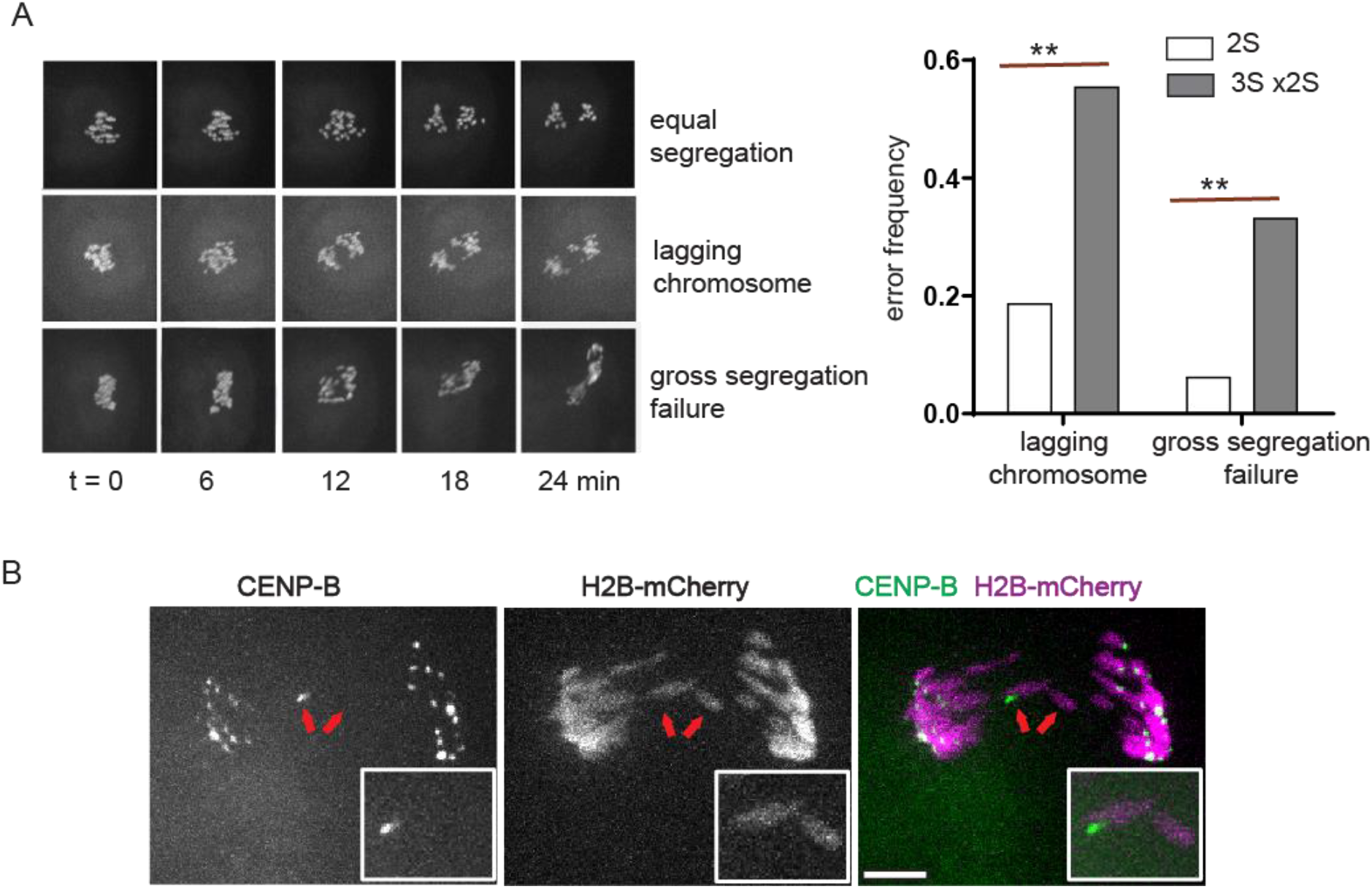
Asymmetric bivalents are associated with meiosis I chromosome segregation errors in 3S × 2S hybrids. A) In oocytes from 3S × 2S hybrids or the 2S control strain, chromosomes were labeled with SiR-DNA for live imaging. Image sequences show examples of equal segregation (top row), lagging chromosomes (middle row), or gross segregation failure (bottom row). Graph shows frequencies of lagging chromosomes or gross segregation failure (n = 72-80 oocytes in each group). **p < 0.01, two-tailed Fisher’s exact test. B) Oocytes from 3S × 2S hybrids expressing CENP-B–EGFP and histone H2B-mCherry were imaged live. Arrows indicate lagging chromosomes. Out of 52 lagging univalents identified from 30 oocytes, 37 had small arrays based on weak or undetectable CENP-B signals (e.g., right arrow). Images are maximum-intensity z-projections. Scale bars 10 μm.

To determine whether lagging chromosomes are a consequence of centromere asymmetry, we repeated the live imaging experiment with expression of CENP-B-GFP to distinguish between larger and smaller array sizes (Figure 3B). Each oocyte contains 20 bivalents at metaphase I, which should resolve into 40 univalents in anaphase I. If lagging occurred at random, univalents with small satellite arrays would account for 5 out of 40 lagging chromosomes (12.5%). Instead, lagging chromosomes were strongly enriched for small satellite arrays (71%, n = 52), indicating that asymmetric bivalents are disproportionately prone to missegregation (p < 10^−20^, exact binomial test). The relatively low frequency of lagging chromosomes in 2S oocytes indicates that small arrays are prone to errors within asymmetric bivalents but not within symmetric bivalents. Some lagging chromosomes with large arrays may also originate from asymmetric bivalents, but our live imaging lacks sufficient temporal resolution to reliably determine their origin.

The live imaging data indicate that centromere asymmetry leads to MI chromosome segregation errors. As an independent measure of errors, we tested for aneuploidy in MII eggs using whole-genome sequencing to determine chromosome copy numbers (Figure 4A). We find increased frequency of aneuploidy in eggs from 3S × 2S hybrid oocytes compared to eggs from 2S control oocytes (Figure 4B). In both cases, loss or gain of a single chromatid is more common than loss or gain of an intact univalent (i.e., copy number of 1 or 3 instead of 0 or 4, Figure 4C), suggesting premature loss of centromere cohesion allowing sister chromatids to separate in meiosis I, consistent with previous studies in mouse oocytes. (Terasa et al., 2010; Treff et al.,2016)

**Figure 4:**
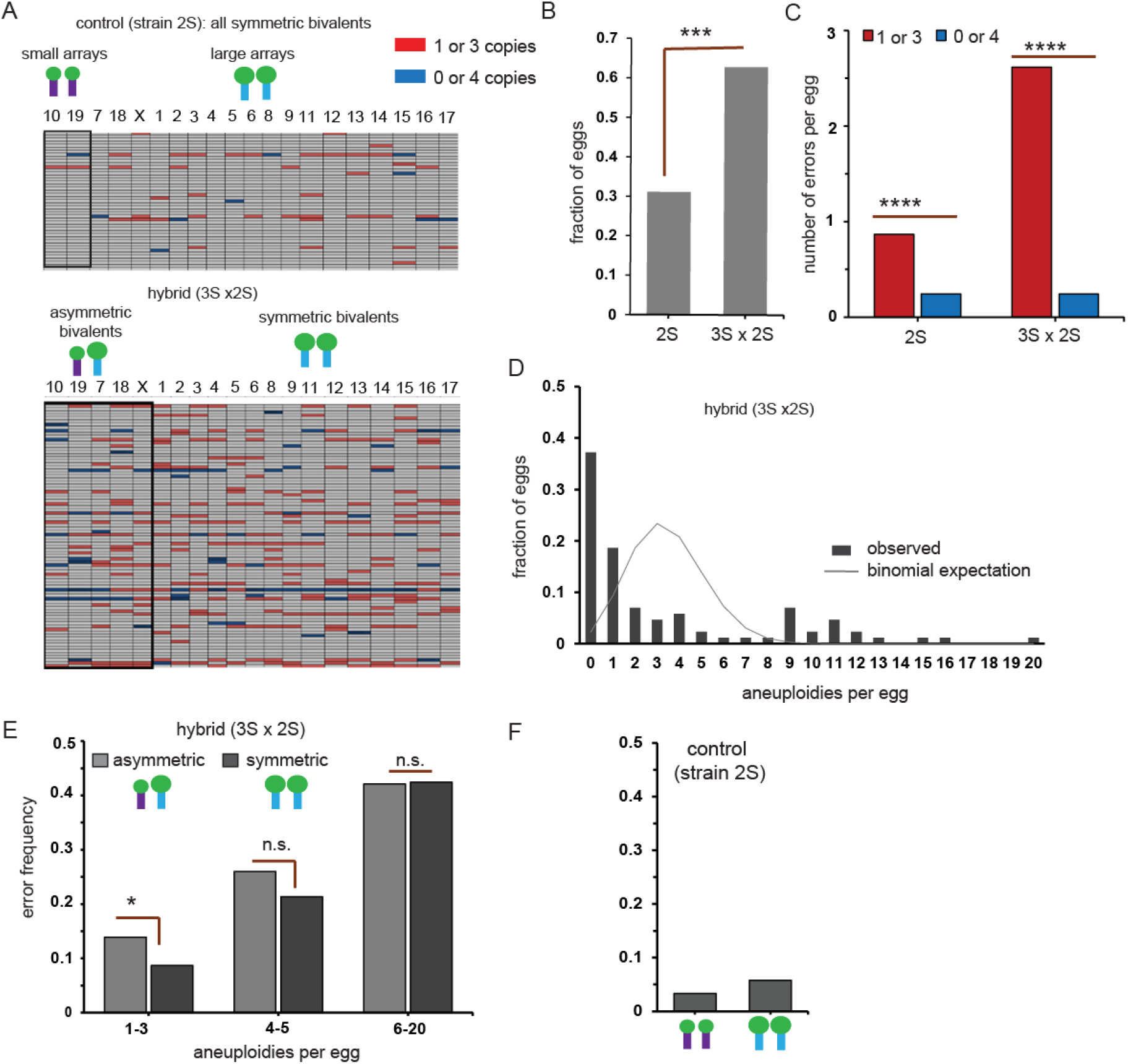
Whole-genome sequencing reveals widespread aneuploidy in 3S × 2S hybrid eggs. Individual MII eggs from 3S × 2S hybrids (n=86) or the 2S control strain (n=45) were processed for single-cell whole genome sequencing to determine chromosome copy numbers. A) Copy-number profiles: each row represents one egg, and each column represents a chromosome. Chromosomes 10 and 19 (first two columns) have small arrays in the 2S strain, and chromosomes 10, 19, 7, 18, and X (first five columns) have asymmetric bivalents in the 3S × 2S hybrid. Colored cells indicate aneuploidy: red for copy number of 1 or 3, or blue for copy number of 0 or 4. B) Fraction of MII eggs containing one or more aneuploidies. ***p = 0.008, two-tailed Fisher’s exact test. C) Number of chromosome copy-number abnormalities per egg, categorized as 1 or 3 copies vs 0 or 4 copies. ****p = 0.0001, two-tailed Fisher’s exact test. D) Distribution of chromosome copy-number abnormalities per egg in 3S × 2S hybrids. Bars show the observed frequency distribution. Line indicates the expected binomial distribution calculated from the observed error rate (0.173) per bivalent in hybrids. E) Data from 3S × 2S hybrids were grouped by number of copy-number abnormalities per egg. Within each group, abnormalities were categorized as coming from asymmetric or symmetric bivalents. *p = 0.045, n.s. not significant, one-tailed Fisher’s exact test. F) For 2S controls, abnormalities were categorized as coming from small or large arrays.

If segregation errors are independent events (i.e., behavior of one bivalent does not affect others in the same cell), the distribution of errors should follow a binomial distribution determined by the overall error frequency per bivalent (0.17 for hybrid oocytes). Our observations from hybrid oocytes are not consistent with this prediction, as euploid eggs and eggs with large numbers of errors are over-represented compared to the binomial distribution (Figure 4D). These findings are consistent with our live imaging data (Figure 3A) indicating two sources of errors. Lagging chromosomes would be expected to generate eggs with relatively small numbers of errors, whereas gross segregation failure would generate eggs with many errors as observed in our aneuploidy data for hybrid oocytes. In eggs with relatively few errors (≤3), we find that error frequencies are higher for asymmetric vs symmetric bivalents (Figure 4E), consistent with our finding that lagging chromosomes frequently arise from asymmetric bivalents, which would be expected to generate eggs with relatively small numbers of errors. Error frequencies are similar, however, for asymmetric and symmetric bivalents in eggs with more errors (≥6), suggesting that these eggs result from gross segregation failure that affects all chromosomes equally. Overall, our aneuploidy measurements indicate that pairing small and large satellite arrays in asymmetric bivalents increases the frequency of errors by both generating lagging chromosomes and disrupting chromosome segregation more broadly. Centromeres with small satellite arrays are not prone to errors in symmetric bivalents (Figure 4F), indicating that array size asymmetry is the key risk factor.

### Functional consequences of asymmetry with large centromere arrays

Our analyses of the 3S × 2S hybrid raise the question of whether the effects of array size asymmetry depend on the presence of extremely small arrays from CHPO within an asymmetric bivalent. To address this, we generated hybrids with PWK, which has the largest documented minor satellite arrays (Arora 2021). We crossed PWK to FVB, which has moderate sized arrays, keeping the ~25-fold difference in satellite array size between homologs comparable to the 3S × 2S hybrid but at the other end of the array size distribution (Figure 5A). Centromeres of homologous chromosomes in PWK × FVB hybrid oocytes exhibited clear differences in CENP-B-GFP signal intensity (Figure 5B), confirming the expected satellite array asymmetry. However, CENP-A levels were similar between centromeres within individual bivalents (Figure 5C), and bivalents were largely aligned near the spindle equator at metaphase I (Figure 5D) in hybrid oocytes with modest differences relative to control FVB oocytes. Chromosome segregation defects were infrequent and comparable to control oocytes based on both live imaging (Figure 6A) and chromosome counting in MII eggs (Figure 6B). Collectively, these results indicate that satellite array asymmetry is not sufficient to induce meiotic defects.

**Figure 5:**
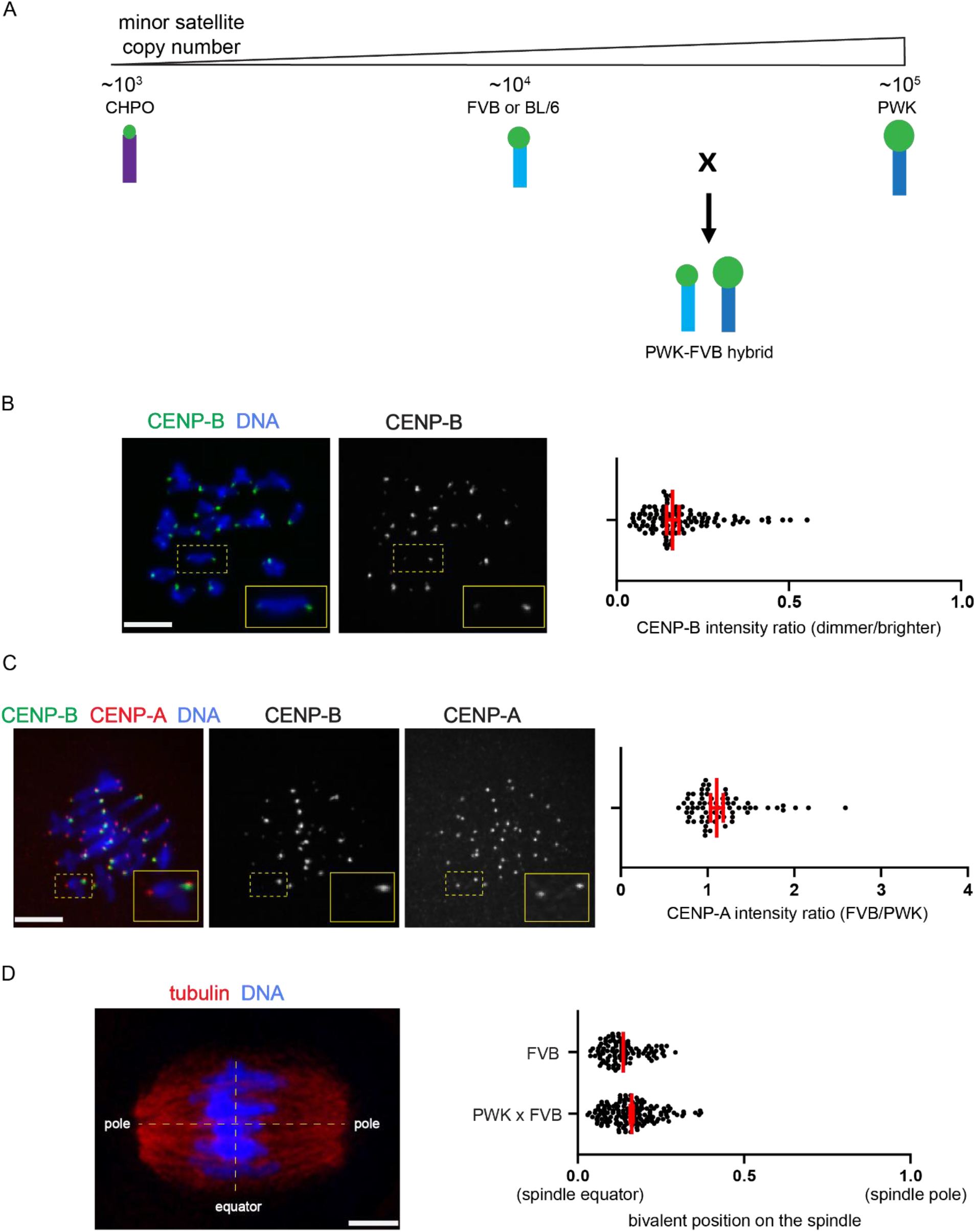
Bivalents in PWK × FVB hybrids exhibit symmetric CENP-A levels and centered position on the MI spindle. A) Generation of PWK × FVB hybrids. B-C) Metaphase I oocytes from PWK × FVB hybrids expressing CENP-B–EGFP. The ratio of CENP-B intensities between centromeres of paired homologous chromosomes was calculated as dimmer/brighter CENP-B signals. Each dot represents a single bivalent; n = 100 pooled from 15 oocytes from 3 independent experiments. D) Metaphase I oocytes from PWK × FVB hybrids expressing CENP-B–EGFP and stained for CENP-A. CENP-B intensities were used to determine small vs large arrays within each bivalent, and the ratio of CENP-A intensities was calculated as small/large (FVB/PWK). Each dot represents a single bivalent; n = 70 pooled from 10 oocytes from 2 independent experiments. E) Metaphase I oocytes from FVB or from PWK × FVB hybrids were fixed and stained for tubulin. Bivalent positions along the spindle axis (horizonal yellow line) were measured relative to the spindle equator (vertical yellow line). Distance from the spindle equator to the pole is defined as 1 for each cell to normalize for variation in spindle size. Each dot represents a single bivalent; n = 100-151 from 10 oocytes from 2 independent experiments. Images are maximum-intensity z-projections. Scale bar 10μm.

**Figure 6:**
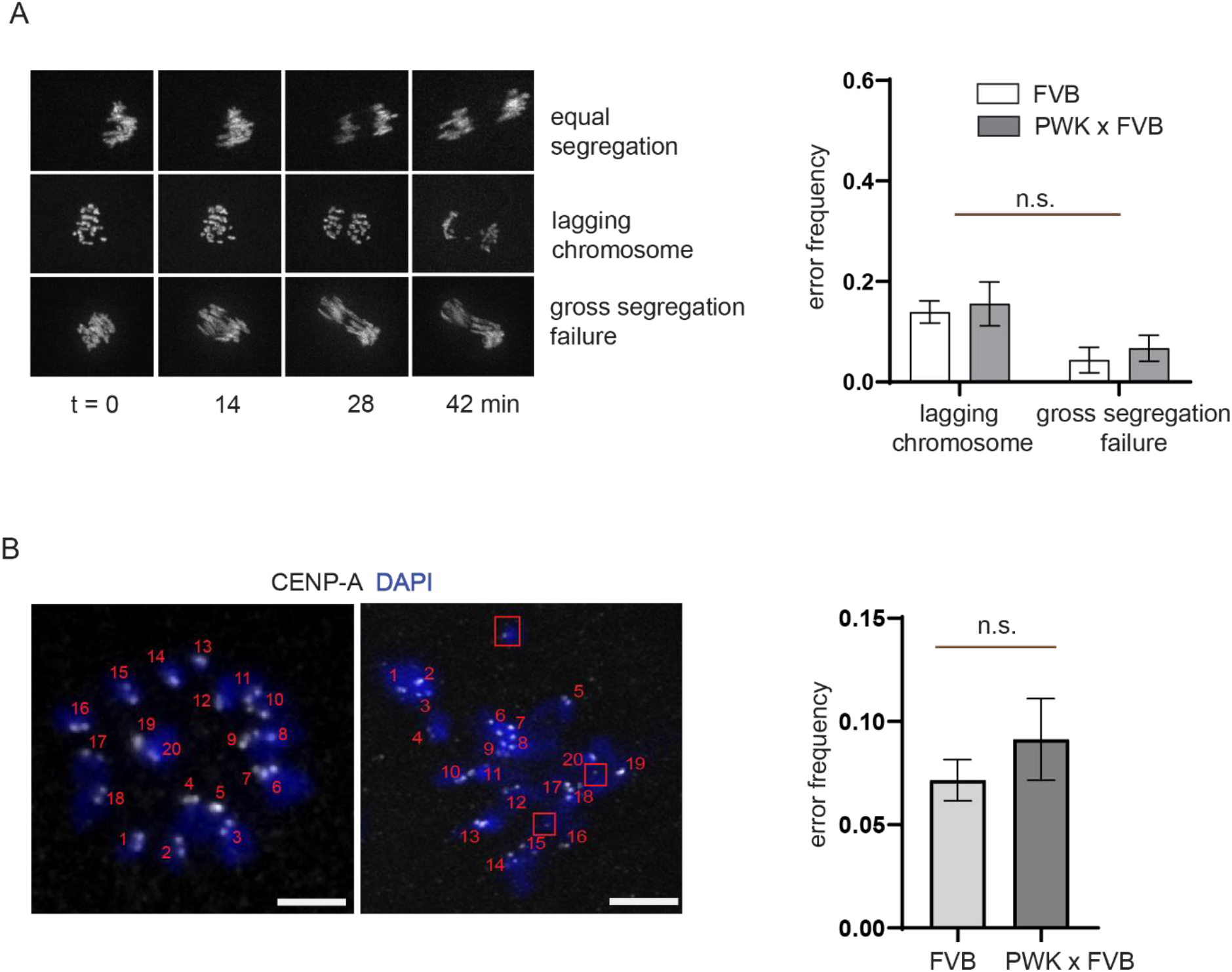
PWK × FVB hybrids show low chromosome segregation error rates during meiosis I. A) Oocytes from FVB or from PWK × FVB hybrids expressing histone H2B-mCherry were imaged live during meiosis I. Image sequences show examples of equal segregation (top row), lagging chromosomes (middle row), or gross segregation failure (bottom row). Graph shows frequencies of lagging chromosomes or gross segregation failure (n = 43-55 oocytes in each group); n.s.: no significant differences between FVB and hybrid, chi-square test. B) Centromere were labeled by CENP-A staining in MII eggs from FVB or PWK × FVB hybrids, and chromosomes were counted to determine aneuploidy; n = 63-66 eggs; Mann– Whitney test, n.s.: not significant. Images are maximum-intensity z-projections. Scale bar 10μm.

## Discussion

Despite extensive documentation of variation in centromeric satellite DNA, its functional consequences have remained unclear. This gap reflects the challenges of developing experimental model systems, as repetitive sequences are difficult to manipulate by genome editing. By leveraging natural variation in satellite array size across inbred mouse strains, we generated hybrid oocytes to directly test the functional impacts of array size asymmetries. We find that differences in satellite array size translate into functional differences at centromeres only when one array is small enough to constrain the amount of centromere chromatin.

We previously showed that CENP-A nucleosomes occupy most of the small minor satellite arrays at CHPO centromeres but only a small fraction of larger arrays. Moreover, the small arrays have reduced levels of CENP-A chromatin and reduced abundance of kinetochore proteins compared to larger arrays (Chmatal et al. 2014, Iwata-Otsubo et al. 2017). These observations suggest that CENP-A nucleosomes are restricted to minor satellite so that small arrays limit the amount of centromere chromatin. Under these conditions, array size asymmetry leads to centromere chromatin and kinetochore asymmetry in meiosis I and functional asymmetry in interactions with spindle microtubules. We now show that this functional asymmetry leads to chromosome segregation errors and aneuploidy, without the complications of meiotic trivalents present in previous CHPO hybrids. In contrast, our findings with the PWK × FVB hybrid indicate that larger arrays do not proportionally expand centromere chromatin, which is symmetric despite large differences in satellite array size, and we do not find evidence of asymmetries in kinetochore function in this hybrid.

Our findings extend the impact of array size asymmetry beyond effects on individual bivalents, as aneuploidy is high for all chromosomes in the 3S × 2S hybrid. This finding is consistent with the view that the spindle operates as a mechanically integrated system (Valdez et al., 2023). For example, kinetochore fibers of adjacent chromosomes can interact dynamically with each other, and steric interactions between chromosomes determine the organization of microtubule bundles in the spindle (Sikirzhytski et al. 2014, Matković et al.2022), suggesting that changes in chromosome position as observed with asymmetric bivalents can impact spindle assembly. In addition, signalling from pole-proximal kinetochores via MPS1 kinase can regulate spindle pole function by phosphorylation of Aurora A kinase (Leça et al., 2025). Thus, asymmetric bivalents positioned near spindle poles could impact the spindle as a whole, leading to segregation errors for all chromosomes. Our findings also suggest that errors are associated with premature loss of centromere cohesion, consistent with prior studies (Gabriel et al., 2011; Kuliev et al., 2011; Tyc et al., 2020). The mechanisms underlying this cohesion loss in the presence of asymmetric bivalents will be an important avenue for future research.

Our findings are consistent with recent analyses of human trisomy 21 showing an association with extreme centromere satellite array size asymmetry between homologous chromosomes in the female parent where the meiotic error occurred (Mastrorosa et al., 2025). Strikingly, these cases of extreme asymmetry include the smallest chromosome 21 array sizes yet described in females. The conclusions from the human study mirror the principle that emerges from our mouse hybrids: small centromere arrays are not inherently defective but increase the risk of segregation errors when paired with a larger array (Figure 7). Our findings thus provide a mechanistic basis for how centromere repeat variation could contribute to human meiotic errors and identify small arrays within asymmetric bivalents as a potential source of meiotic vulnerability.

**Figure 7:**
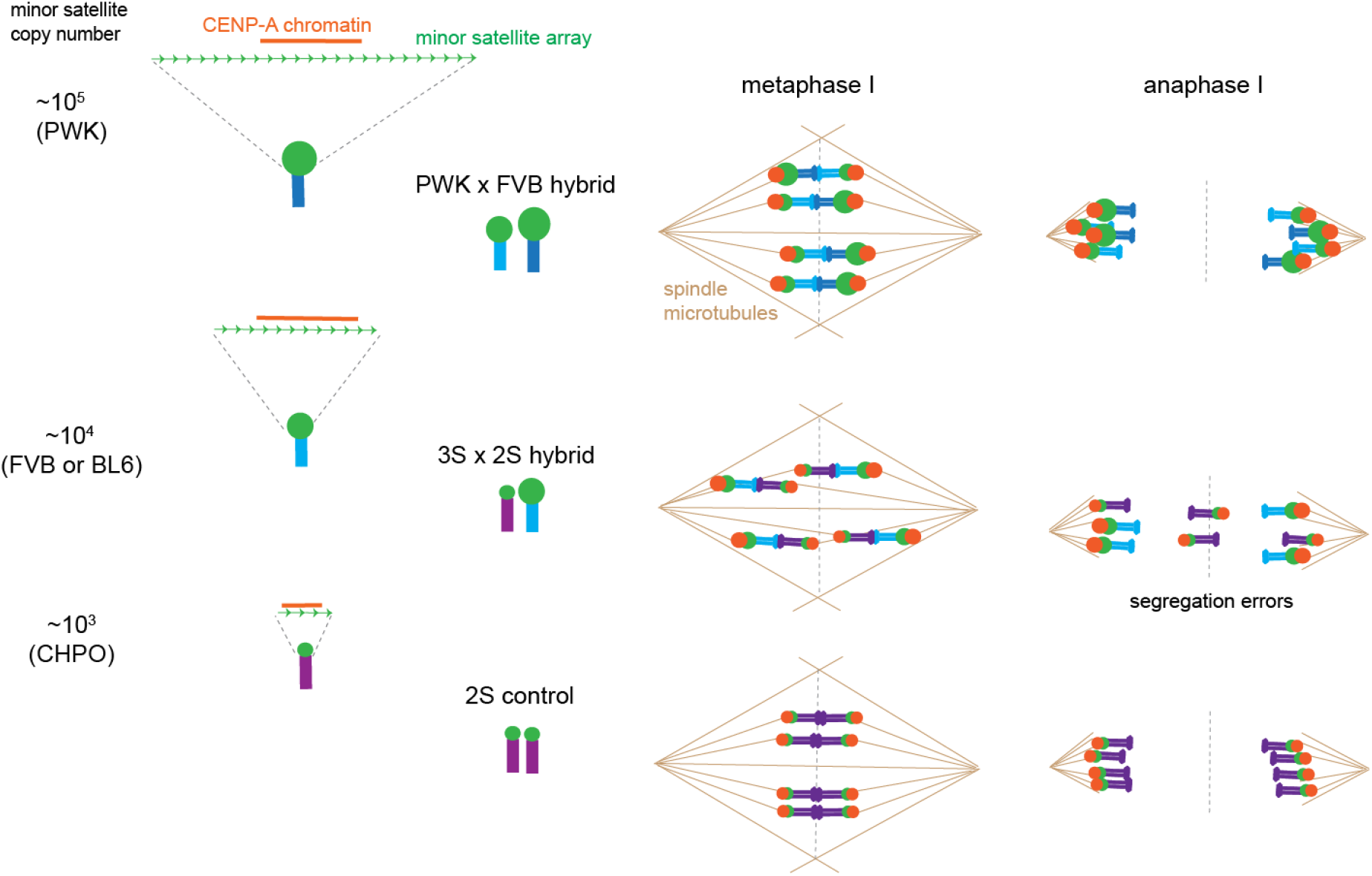
Functional consequences of centromere satellite array size asymmetry. CENP-A chromatin is constrained by small satellite arrays (e.g., CHPO centromeres) but does not scale with larger arrays (FVB, BL6, PWK). Meiotic asymmetries in CENP-A chromatin are observed only in hybrids with small arrays, leading to asymmetric interactions with spindle microtubules in metaphase I and segregation errors in anaphase I.

## Materials and methods

### Mice

Mouse strains were purchased from the Jackson Laboratory (ZALENDE/EiJ, stock #001392 corresponds to CHPO; C57BL/6J, stock# 000664; FVB, stock#001800; PWK, stock# 003715). CHPO/ZALENDE and C57BL/6 (BL6) mice were crossed to generate CHPO hybrids, which were backcrossed to BL6. SNP genotyping was performed using the MiniMUGA mouse genotyping array (Neogen) to identify CHPO or BL/6 centromeres after each backcross (Sigmon et al., 2020). PWK and FVB mice were crossed to generate PWK × FVB hybrids. All mice used in this study were female at 7–14 weeks old. All animal experiments were approved by the Institutional Animal Care and Use Committee of the University of Pennsylvania and were consistent with the National Institutes of Health guidelines (protocol: #804882).

### Oocyte collection and culture

Fully grown germinal vesicle (GV)–stage oocytes were recovered in bicarbonate-free MEM supplemented with HEPES and polyvinylpyrrolidone (MEM–PVP) and mechanically cleared of cumulus cells. Oocytes were cultured in Chatot–Ziomek–Bavister (CZB) medium containing 2.5 mM milrinone under mineral oil (Sigma) at 37 °C in a 5% CO_2_ environment to maintain prophase I arrest. Meiotic resumption was initiated by transferring oocytes through successive washes into milrinone-free CZB medium.

### Immunostaining of MI oocytes

At 7-10 h following milrinone washout, oocytes were fixed in 2% paraformaldehyde in PBS containing 0.1% Triton X-100 for 20 min. For detection of CENP-A, samples were incubated with λ-phosphatase (80 U/mL; New England Biolabs) for 2 h at 30 °C, permeabilized in 0.1% Triton X-100 in PBS for 15 min at room temperature, and then incubated in blocking solution (0.1% BSA and 0.01% Tween-20 in PBS) for 15 min. Primary antibody incubation was carried out using rabbit anti–CENP-A (1:200; Cell Signaling Technology, #2048S) for 1 h, followed by incubation with fluorescently conjugated secondary antibodies (goat or donkey anti-rabbit IgG, 1:500; or anti-human IgG, 1:100; Life Technologies) for 1 h. Between each step, oocytes were washed three times in blocking buffer for 10–15 min per wash. Samples were mounted in Vectashield medium containing DAPI. Image analysis was performed in Fiji/ImageJ. Z-slices encompassing individual bivalents were combined into sum projections, and centromeres were manually outlined. Integrated fluorescence intensity was measured for each centromere following background subtraction, using a nearby region to estimate background signal.

### Oocyte microinjection and live imaging

To visualize chromosome dynamics by live imaging during meiosis I, oocytes were incubated with 250 nM of the far-red DNA dye SiR-DNA. In some experiments, centromeres were also labeled by expression of CENP-B–EGFP, allowing discrimination of centromeres with differing satellite array sizes during live imaging. CENP-B–EGFP cRNA was synthesized using the TranscriptAid T7 High Yield Transcription Kit, purified byMEGAclear™ Transcription Clean-Up Kit, and microinjected at a concentration of 200 ng/ul. Meiotic resumption was induced by milrinone washout, and oocytes were transferred to CZB medium under mineral oil in glass-bottom dishes (FluoroDish FD35-100). Imaging was performed in a heated environmental chamber with a stage-top incubator (PeCon) maintained at 37 °C and 5% CO_2_. Confocal imaging was carried out on a Leica DMI4000 B microscope equipped with a 63× 1.3 NA glycerol-immersion objective, a piezo-driven Z stage (Applied Scientific Instrumentation), and a Yokogawa spinning-disk confocal scanner. Images were acquired using an EM-CCD camera (Hamamatsu ImageEM C9100-13) with 488- and 593-nm lasers controlled by MetaMorph software. For analysis of chromosome segregation, oocytes were imaged 7-10 hours after GVBD until anaphase I.

### Bivalent positioning assay

For analysis of bivalent positioning on the MI spindle, GV-stage oocytes were collected and allowed to mature in vitro to metaphase I (7 –10 h after GVBD). Oocytes were then fixed and stained for α-tubulin and DNA. The spindle axis was defined by the two poles visualized by tubulin staining, and the position each bivalent was projected onto this axis and calculated relative to the nearest pole and the spindle midzone, defined as the midpoint between the two spindle poles.

### Centromere counting assay in MII eggs

GV-stage oocytes were collected from hormonally stimulated females and cultured in milrinone-containing medium to maintain prophase I arrest. Meiotic resumption was induced by washing out milrinone, and oocytes were cultured to MII. MII eggs were then incubated in CZB medium containing 100 µM monastrol (kinesin-5 inhibitor) for 1.5 h at 37 °C in a humidified atmosphere of 5% CO_2_. Eggs were fixed in 2% paraformaldehyde, stained for centromeres using anti–CENP-A, and mounted in Vectashield medium containing DAPI. Z-stack images were acquired at 0.3-µm intervals using the confocal imaging setup described above. Centromere number per egg was determined by manual counting of CENP-A foci using Fiji/ImageJ.

### Whole Genome Sequencing Analysis

To assess the impact of centromeric asymmetry on chromosome segregation fidelity, we performed single-cell whole genome sequencing on metaphase II eggs from each mouse line. GV oocytes were collected from PMSG primed females and allow to progress to MII egg for 16hours in Milrinone-free CZB medium. Zona pellucida was then removed by Acid Tyrode’s solution treatment and polar body were mechanically stripped away by passing through small-pore micropipette. Only MII eggs with visible polar body were used in experiments and polar body removal was confirmed under dissecting microscope. Single Cell whole genome amplification was performed using the Sigma-Aldrich GenomePlex WGA4 kit. Whole genome libraries were constructed by using Illumina DNA PCR-Free Prep kit. Libraries were pooled and sequenced on an Illumina MiSeq platform in single-end 100 bp mode, targeting ~0.1–0.4× genome coverage per cell depending on pooling density. This sequencing depth yielded approximately 10,000 to 40,000 reads per chromosome, sufficient to detect full chromosome copy number changes. Sequencing adapters were trimmed using the Trim Galore! Tool (Galaxy bioinformatics suite). Quality-trimmed reads were aligned to the Mus musculus mm10 reference genome using Bowtie2, generating BAM files. AneuFinder (development version, https://bioconductor.org/packages/release/bioc/html/AneuFinder.html) was then used to detect, analyze copy number variations in single-cell whole-genome libraries.

